# Assessment of Circulating Copy Number Variant Detection for Cancer Screening

**DOI:** 10.1101/041848

**Authors:** Bhuvan Molparia, Eshaan Nichani, Ali Torkamani

## Abstract

Current high-sensitivity cancer screening methods suffer from false positive rates that lead to numerous unnecessary procedures and questionable public health benefit overall. Detection of circulating tumor DNA (ctDNA) has the potential to transform cancer screening. Thus far, nearly all ctDNA studies have focused on detection of tumor-specific point mutations. However, ctDNA point mutation detection methods developed to date lack either the scope or sensitivity necessary to be useful for cancer screening, due to the extremely low (<1%) ctDNA fraction derived from early stage tumors. We suggest that tumor-derived copy number variant (CNV) detection is theoretically a superior means of ctDNA-based cancer screening for many tumor types, given that, relative to point mutations, each individual tumor CNV contributes a much larger number of ctDNA fragments to the overall pool of circulating DNA. Here we perform an in silico assessment of the potential for ctDNA CNV-based cancer screening across many common cancers.

## INTRODUCTION

According to the National Cancer Institute, 5 year survival rates of cancer patients are the highest when cancer is detected and treated at an early, localized, stage. Currently, there are a number of different cancer-type specific biomarkers used to detect cancer at an early stage; however most of them are associated with alarmingly high false positive rates (FPRs). For example, ovarian cancer screening using the CA-125 biomarker (1) along with transvaginal ultrasonography has a sensitivity of ~90% but a FPR of 57% (2). Mammography for breast cancer screening has a FPR of 40-60% over 10 years of screening (3), Cologuard^®^ for colorectal cancer screen has a FPR of 13.4% (4), and PSA for prostate cancer screening has a FPR of 20-30% when the test aims to detect >80% of cancers (5). False positive results, and sometimes screening methods themselves, tend to lead to invasive and uncomfortable procedures that are associated with risk to otherwise healthy individuals; e.g. radiation exposure during mammography and surgery or biopsy in the case of other tumor types. These unnecessary procedures, unfortunately, lead to adverse events in approximately 15% of cases (6). High false positive rates along with high adverse event rates for follow-up procedures place a significant proportion of the healthy population at unnecessary risk. Thus, an alternative and highly accurate non-invasive method for early cancer detection, especially a global test for multiple types of cancer, would both reduce the rate and impact of false positive results on otherwise healthy individuals, and could lead to substantial improvements in survival and quality of life of cancer patients.

Copy number variations (CNVs), like point mutations, are common and causal for a large proportion of cancer types (7, 8). CNV based classification of tumor subtypes has been demonstrated previously, though these methods have been focused on gene level events and the stratification of tumors from a single organ system into clinically relevant subtypes - rather than assignment of a CNV profile to a tissue of origin (9-11). The recent development of non-invasive prenatal tests (NIPT), which involve whole-genome shotgun sequencing of circulating DNA (100-200bp fragments) (12) in pregnant women in order to detect fetal chromosomal abnormalities, has revealed the potential of circulating DNA tests for the detection of cancer (13). However, the tumors incidentally detected by NIPT testing to date have been relatively bulky with large and numerous chromosome scale abnormalities, or were simply hematological malignancies, and thus not representative of the types of tumors that should ideally be detected by routine cancer screening.

A major obstacle to the utilization of circulating DNA tests in cancer screening, both for small variant as well as CNV detection is the low fraction of circulating tumor-derived DNA (ctDNA). NIPT testing can detect major chromosomal abnormalities in the fetus, in a context where 10-50% (14, 15) of the circulating DNA in a pregnant woman’s bloodstream is derived from the fetus. On the other hand, the fraction of ctDNA in the bloodstream is <1%, except for in circumstances where a screening test is irrelevant i.e. large and widely disseminated tumors (16). Given that the weak ctDNA mutation signature is dispersed over a large genomic region in the case of tumor-derived CNVs, we reasoned that detection of large circulating tumor-derived CNVs via sequencing may be a more viable cancer screening application of ctDNA sequencing, relative to ctDNA point mutation detection methodologies which either have modest sensitivity or must be customized to a patient and thus are only applicable to cancer treatment monitoring (17-19). Moreover, ctDNA CNV signatures are more likely to be useful for determining the tissue of origin of a tumor, a characteristic that is important for clinical follow-up in a cancer screening setting.

In this light, we explore the potential for ctDNA CNV detection for cancer screening by evaluating the ability to detect and differentiate tumor types via large tumor CNV events (5 megabases or greater) that are theoretically detectable via ctDNA sequencing (15). We demonstrate that, for many tumor types, including those not necessarily enriched with CNV events, it is theoretically possible to accurately detect and classify tumor types via large ctDNA detectable tumor-derived CNVs.

## RESULTS

### Sample Clustering Using SAX Representation

First, we divided the human genome into segments of sizes 5, 6, 7.5, 10, 30, 40, 60, 75, and 100 megabases – segment sizes representing a range of tumor-derived CNV event sizes (7) that are hypothetically detectable via ctDNA sequencing at low sequence depth (15). We then overlaid the CNV profile of each tumor sample across these bins and determined the copy number variant status of each bin based on the raw segment duplication values. Symbolic Aggregate Approximation, a method developed to convert time series data to a representation amenable to clustering, classification, and matching algorithms (20), was selected for this purpose. SAX transformation reduces the dimensionality of tumor CNV profiles, from their continuous numerical representation with relatively precisely defined CNV boundaries, to a symbolic representation robust to variability in somatic CNV boundaries across samples, and amenable to both simple and complex classification algorithms. This simplifies the overall CNV profile of a tumor while allowing for the identification of “critical” regions for CNV based tumor classification. Specifically, we transformed each segment to a SAX representation with cardinality 5, representing normal or 2 copies, 1 copy amplification, more than 2 copy amplifications, 1 copy deletion and 2 copy deletion of the genomic segment. This process was meant to simulate the detection of CNVs from ctDNA by read counting applications applied across large genomic segments (21, 22). 11 major pan-cancer (10) solid tumor types were considered: breast adenocarcinoma (BRCA), lung adenocarcinoma (LUAD), lung squamous cell carcinoma (LUSC), uterine corpus endometrial carcinoma (UCEC), glioblastoma multiforme (GBM), head and neck squamous cell carcinoma (HNSC), colon and rectal carcinoma (COAD, READ), bladder urothelial carcinoma (BLCA), kidney renal clear cell carcinoma (KIRC), ovarian serous carcinoma (OV).

We performed unsupervised clustering using the SAX transformed CNV values, at each segment size, to determine whether simple clustering would be sufficient to differentiate different tumor types from one another and from normal samples. Unsupervised clustering utilizing an unfiltered set of all genomic segments did not effectively separate tumor types from one another (not shown), therefore, we performed restricted clustering to genomic segments deemed relevant for distinguishing tumor types in later applied random forest classification model (*see below*). The resulting heat maps demonstrate some degree of separation of disparate tumor types from one another, but clearly demonstrates that, at CNV sizes detectable by low pass ctDNA sequencing methods; simple clustering does not sufficiently separate tumor types from one another for clinical classification applications (Figure 1 and Supplemental Figures). Certain tumor types like GBM and KIRC form cohesive, though not complete, blocks of clustered samples, while most others demonstrated a tendency to form close but intermixed clusters with other tumor types. These results suggest that classification of tumor type by CNV profile is possible, though requires more sophisticated methodology to account for heterogeneity within tumor types and similarity across tumor types (10).

**Figure 1:**
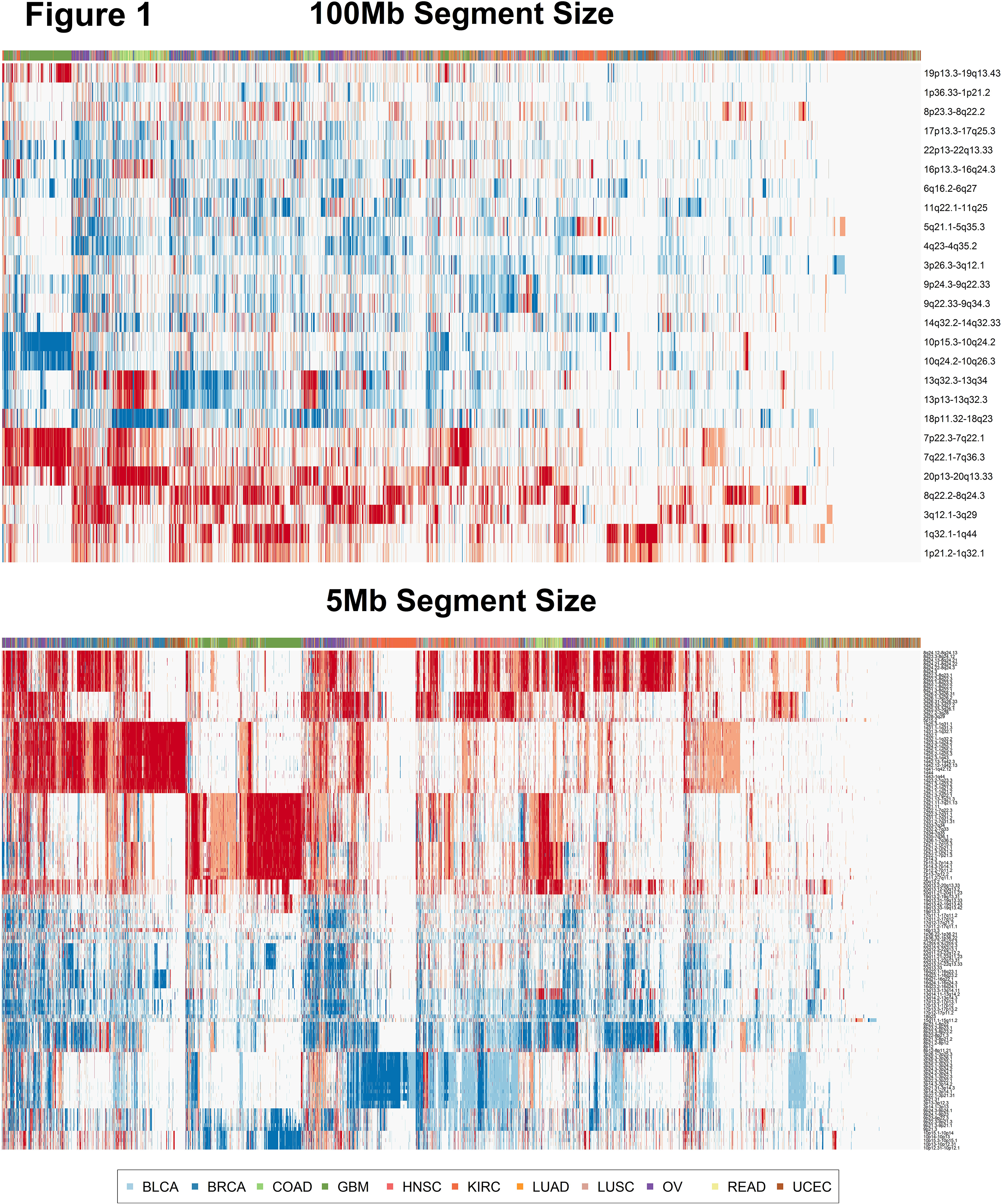
Sample Clustering with Symbolic Aggregate Approximation (SAX) Representation. Heat maps showing the results of unsupervised clustering using SAX transformed data at the 100 Mb segment size (top panel) and 5 Mb segment size (bottom panel). Deletions are colored blue and amplifications are colored red. We achieve some extent of separation of disparate tumor types from one another, with some cancers like KIRC and GBM forming compact but incomplete groups while most others remain in intermixed clusters.

### Nearest Neighbor Classification

Given the promising but limited success of unsupervised clustering for cancer type classification, we sought to determine whether cancer type can be determined by a k nearest neighbor approach, which should account for enrichment of cancer types within intermixed clusters. This model was readily capable of distinguishing cancer samples from normal samples with a true positive rate (TPR) of 0.80 and a positive predictive value (PPV) of 0.9995 at the 100 Mb segment size threshold; suggesting, perhaps unsurprisingly that, in theory, a ctDNA CNV profile, even at coarse resolution, can differentiate between healthy individuals and those with cancer with a negligible false positive rate. The TPR and PPV were stable across all segment sizes implying that the major differentiating factor between cancer and normal samples are relatively large scale genomic copy number changes and high resolution of CNVs is not required to differentiate samples from healthy and cancer patients (Supplemental Table 1). Thus, simple detection of cancer via ctDNA sequencing and CNV detection is straightforward. However, certain CNV poor cancer types, such as pancreatic adenocarcinomas, prostate adenocarcinomas and thyroid carcinomas are difficult to distinguish from normal samples given their flat CNV profiles overall.

When the nearest neighbor approach was utilized in attempt to determine cancer type or origin, an overall accuracy of 0.691 was observed at the 100Mb segment size. At the 5 Mb segment size the overall accuracy of cancer type prediction was 0.694 - a non-significant improvement over the accuracy at 100Mb segment size (Supplemental Table 2). Again, most of the poor performers were cancer types with flat CNV profiles while kidney cancers and GBM performed the best (Table 1). This is consistent with the finding that KIRC and GBM also formed more or less consistent clusters during simple clustering. Thus, while the nearest neighbor approach readily distinguished cancer from normal profiles, at CNV resolution detectable by ctDNA sequencing, accuracy remains insufficient for differentiation of tumor types from one another.

**Table 1:** Performance of the KNN and random forest models in determining cancer type. The predictive performance of the KNN model and the optimal performance of the random forest model-based on the point on the ROC curve nearest to 100% specificity and sensitivity - per cancer type are listed at the 100 Mb and 5 Mb segment size thresholds. PPV: Positive predictive value; TPR: True positive rate.

| Sample Type | KNN |  |  |  | Random Forest (Optimal) |  |  |  |
| --- | --- | --- | --- | --- | --- | --- | --- | --- |
|  | 100 Mb |  | 5 Mb |  | 100 Mb |  | 5 Mb |  |
|  | TPR | PPV | TPR | PPV | TPR | PPV | TPR | PPV |
| Thyroid carcinoma | 0.088 | 0.182 | 0.105 | 0.245 | 0.200 | 0.521 | 0.291 | 0.558 |
| Pancreatic adenocarcinoma | 0.000 | 0.000 | 0.020 | 0.500 | 0.486 | 0.610 | 0.627 | 0.696 |
| Uterine Corpus Endometrial Carcinoma | 0.252 | 0.284 | 0.325 | 0.221 | 0.669 | 0.709 | 0.684 | 0.730 |
| Stomach adenocarcinoma | 0.357 | 0.303 | 0.232 | 0.277 | 0.758 | 0.777 | 0.817 | 0.824 |
| Prostate adenocarcinoma | 0.130 | 0.213 | 0.206 | 0.415 | 0.550 | 0.669 | 0.824 | 0.834 |
| Colon adenocarcinoma | 0.673 | 0.536 | 0.645 | 0.477 | 0.830 | 0.837 | 0.834 | 0.843 |
| Bladder Urothelial Carcinoma | 0.115 | 0.429 | 0.077 | 0.500 | 0.810 | 0.808 | 0.839 | 0.845 |
| Kidney renal papillary cell carcinoma | 0.574 | 0.684 | 0.618 | 0.627 | 0.875 | 0.881 | 0.841 | 0.848 |
| Lung adenocarcinoma | 0.113 | 0.536 | 0.135 | 0.400 | 0.822 | 0.830 | 0.853 | 0.857 |
| Esophageal carcinoma | 0.022 | 1.000 | 0.000 | NA | 0.795 | 0.805 | 0.854 | 0.853 |
| Adrenocortical carcinoma | 0.409 | 0.750 | 0.409 | 0.692 | 0.911 | 0.909 | 0.856 | 0.864 |
| Head and Neck squamous cell carcinoma | 0.280 | 0.298 | 0.333 | 0.352 | 0.834 | 0.840 | 0.872 | 0.876 |
| Cervical squamous cell carcinoma and endocervical adenocarcinoma | 0.042 | 0.250 | 0.097 | 0.318 | 0.855 | 0.858 | 0.875 | 0.876 |
| Kidney Chromophobe | 0.882 | 0.789 | 0.882 | 0.682 | 0.879 | 0.888 | 0.879 | 0.889 |
| Breast invasive carcinoma | 0.665 | 0.365 | 0.482 | 0.519 | 0.862 | 0.865 | 0.881 | 0.883 |
| Liver hepatocellular carcinoma | 0.150 | 0.750 | 0.390 | 0.513 | 0.844 | 0.851 | 0.884 | 0.884 |
| Rectum adenocarcinoma | 0.027 | 0.250 | 0.000 | 0.000 | 0.916 | 0.916 | 0.886 | 0.889 |
| Uterine Carcinosarcoma | 0.000 | NA | 0.000 | NA | 0.893 | 0.886 | 0.893 | 0.885 |
| Skin Cutaneous Melanoma | 0.212 | 0.833 | 0.297 | 0.833 | 0.847 | 0.853 | 0.900 | 0.902 |
| Kidney renal clear cell carcinoma | 0.746 | 0.585 | 0.754 | 0.537 | 0.881 | 0.888 | 0.902 | 0.907 |
| Lung squamous cell carcinoma | 0.488 | 0.438 | 0.545 | 0.493 | 0.860 | 0.868 | 0.910 | 0.910 |
| Brain Lower Grade Glioma | 0.478 | 0.637 | 0.463 | 0.441 | 0.865 | 0.864 | 0.911 | 0.913 |
| Pheochromocytoma and Paraganglioma | 0.435 | 0.833 | 0.391 | 0.857 | 0.922 | 0.923 | 0.911 | 0.914 |
| Glioblastoma multiforme | 0.783 | 0.675 | 0.811 | 0.577 | 0.929 | 0.930 | 0.944 | 0.944 |
| Ovarian serous cystadenocarcinoma | 0.321 | 0.863 | 0.372 | 0.879 | 0.937 | 0.937 | 0.957 | 0.956 |
| Normal | 1.000 | 0.836 | 0.999 | 0.832 | 0.994 | 0.993 | 0.983 | 0.982 |

### Random Forest Classification

Finally, we utilized a random forest classification model to simulate the (near) optimal classification performance of the ctDNA biomarker (23). The model had an overall accuracy of 0.78 and 0.74 when classifying all tumor types at segment sizes of 5Mb and 100Mb respectively. ROC curves at 5Mb and 100Mb segment size are plotted in Figure 2 for the 11 major solid tumor types, and demonstrate significant differences in performance across tumor types. Additional ROC curves for all thresholds and cancer types can be found in the *Supplemental Data*. Figure 2 demonstrates an overall improvement in the predictive power across most cancer types at the greater segment size resolution, but once again the improvement when increasing resolution from 100Mb to 5Mb segments is not dramatic. Certain cancer types, such as OV, BRCA, GBM and KIRC are consistently and accurately (~95%) assigned to the correct tumor type across all segment sizes. While others, apparently those of squamous histology, such as HNSC, LUSC, and BLCA show a considerable improvement (~5% increase in accuracy) in predictive value as segment size detection resolution is improved. Thus, while cancer profiles can be readily distinguished from normal profiles, determination of the tissue of origin shows variability in performance across tumor types and segment size resolution.

**Figure 2:**
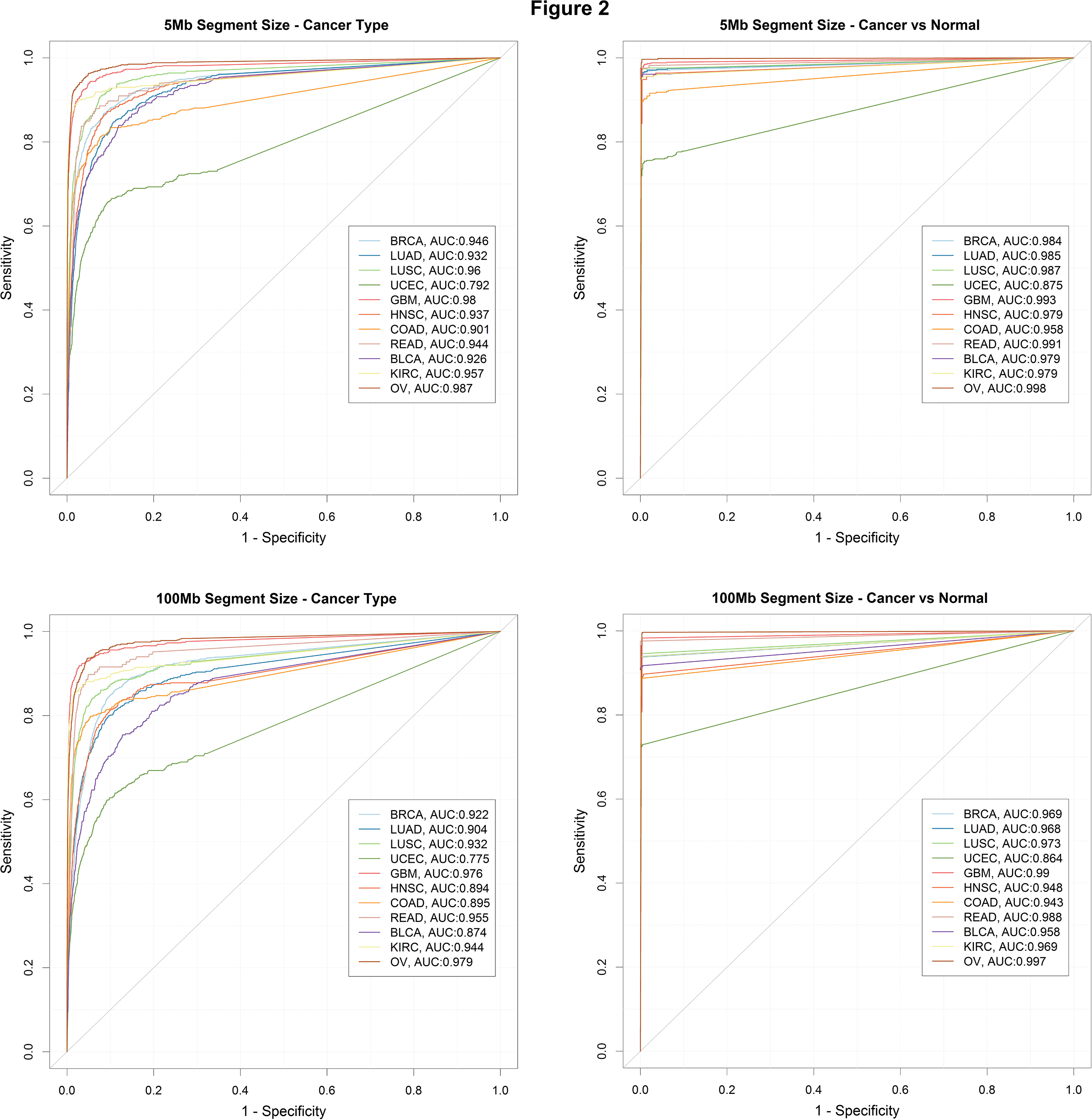
Tumor Classification Performance. ROC curves at 5Mb (top panels) and 100Mb segment size (bottom panels) showing performance of classification of cancer types (left panels)) and cancer vs normal (right panels) for 11 major types of solid tumors - breast adenocarcinoma (BRCA), lung adenocarcinoma (LUAD), lung squamous cell carcinoma (LUSC), uterine corpus endometrial carcinoma (UCEC), glioblastoma multiforme (GBM), head and neck squamous cell carcinoma (HNSC), colon and rectal carcinoma (COAD, READ), bladder urothelial carcinoma (BLCA), kidney renal clear cell carcinoma (KIRC), ovarian serous carcinoma (OV). It can be seen that there is an overall increase in the AUC values when going from a 100 Mb segment size to 5 Mb segment size in both Cancer vs Normal and Cancer Type prediction. The increase in AUC, however, is not dramatic and reflects the robustness of this method at various size resolutions.

The optimal performance for the different cancer types, based on the point on the ROC curve nearest to 100% specificity and sensitivity is presented in Table 1 and Supplemental Table 3. Most cancer types demonstrate a PPV of >80% at the lowest segment size resolution (100Mb) outpacing the PPV of diagnostic tests currently used in clinical practice. OV and GBM demonstrate the best performance (>90% TPR and PPV) suggesting they are excellent candidates for ctDNA CNV based screening applications. Many other tumor types, especially BRCA, UCEC, READ, KIRC, and LUSC also demonstrate excellent TPR and PPV rates (Table 1).

### Tumor Misclassification

Given that our prediction model does not correctly classify all the tumor samples, we investigated whether misclassifications were driven by biological relationships or were true classification errors. The misclassification heat map at the 5 Mb segment size threshold is displayed in Figure 3, additional heat maps for other size thresholds are available as *Supplementary Data*. Many misclassifications were true errors - cancer samples being classified as normal samples (23.4% of errors) or as Breast invasive carcinomas (12.8% of errors). These errors are largely derived from cancer types such as Thyroid Carcinomas that have very poor performance overall and have flat CNV profiles that are difficult to distinguish from normal samples (Supplemental Table 4). Breast cancers tend to be diverse with respect to their cell type of origin and contain molecularly distinct subtypes (10), and thus may mimic CNV profiles of other tumor types. While breast cancer samples themselves were classified accurately, many errors were derived from other tumor types being classified as breast cancer. Given that breast cancer was the largest sample set overall, the imbalance of tumor samples per type is potentially driving this misclassification bias.

**Figure 3:**
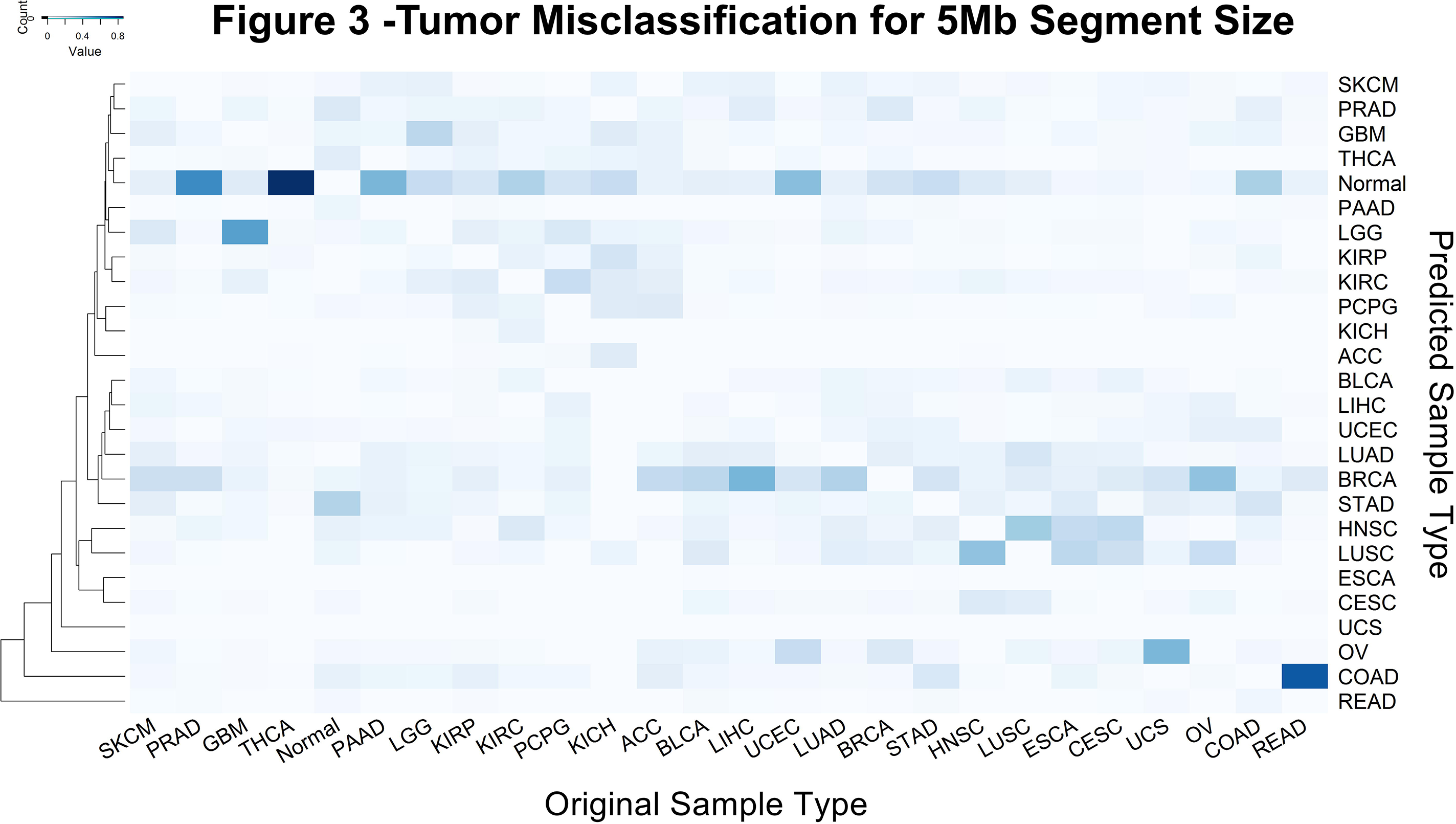
Heat map cluster of tumor types by misclassification frequency. Every column has one cancer type and each row displays the misclassification frequency of that cancer type to the one represented by the row. Correct classifications are set to 0 to highlight the misclassifications. Color indicates the frequency of misclassification with darker color being a higher frequency. Cancers seem to cluster according to their cells of origin and most misclassifications are to cancers with similar cell type. Cancers like THCA, PAAD and PRAD that have low CNV burden are the hardest to detect from Normal samples.

For cancer types without flat CNV profiles, misclassification tended to cluster based on tissue of origin or molecular subtype. For example, squamous cell cancers like LUSC, HNSC, ESCA and CESC show similar misclassification patterns and are often misclassified for one another (Figure 3). Tumors of the gastrointestinal tract, STAD, COAD, and READ are also often misclassified for one another. Similarly, brain cancers GBM and LGG show a high degree of cross misclassification. Cancers originating from the kidneys (KIRC, KIRP, and KICH) form a misclassification cluster with ACC and PCPG. These tissues originate from the intermediate mesoderm, which likely explains their cross misclassifications. Finally, tumors driven by similar genetic mechanisms, e.g. OV and BRCA, were often misclassified for one another. These results suggest that misclassification is often biologically driven and follow clinically addressable patterns.

### Theoretical Power of Detecting Segment CNV

Finally, we performed an in silico assessment of the potential for ctDNA CNV detection at low ctDNA fractions. The utility of the prediction accuracy described herein would only be borne out if ctDNA CNVs are detectable at the given segment size thresholds. Obviously, massively high read depth could potentially be used to detect small ctDNA derived CNVs at low ctDNA fractions (~0.1%). However, we aimed to determine whether predictive accuracy could be balanced with cost and implementation feasibility. To assess the feasibility of ctDNA CNV detection for cancer screening, we estimated the theoretical sequencing depth required to detect a single copy change (p-value < 0.01) at various segment size thresholds and ctDNA fractions. Figure 4 displays the results of this calculation with variance calculated using a Negative Binomial distribution and an artificially high assumed variance of 2 times the mean. At 0.1% ctDNA fraction of total circulating DNA, a 100Mb copy number variant aberration is easily detectable with less than 100 million reads - suggesting that the predictive performance presented herein is technically and clinically achievable in practice.

**Figure 4:**
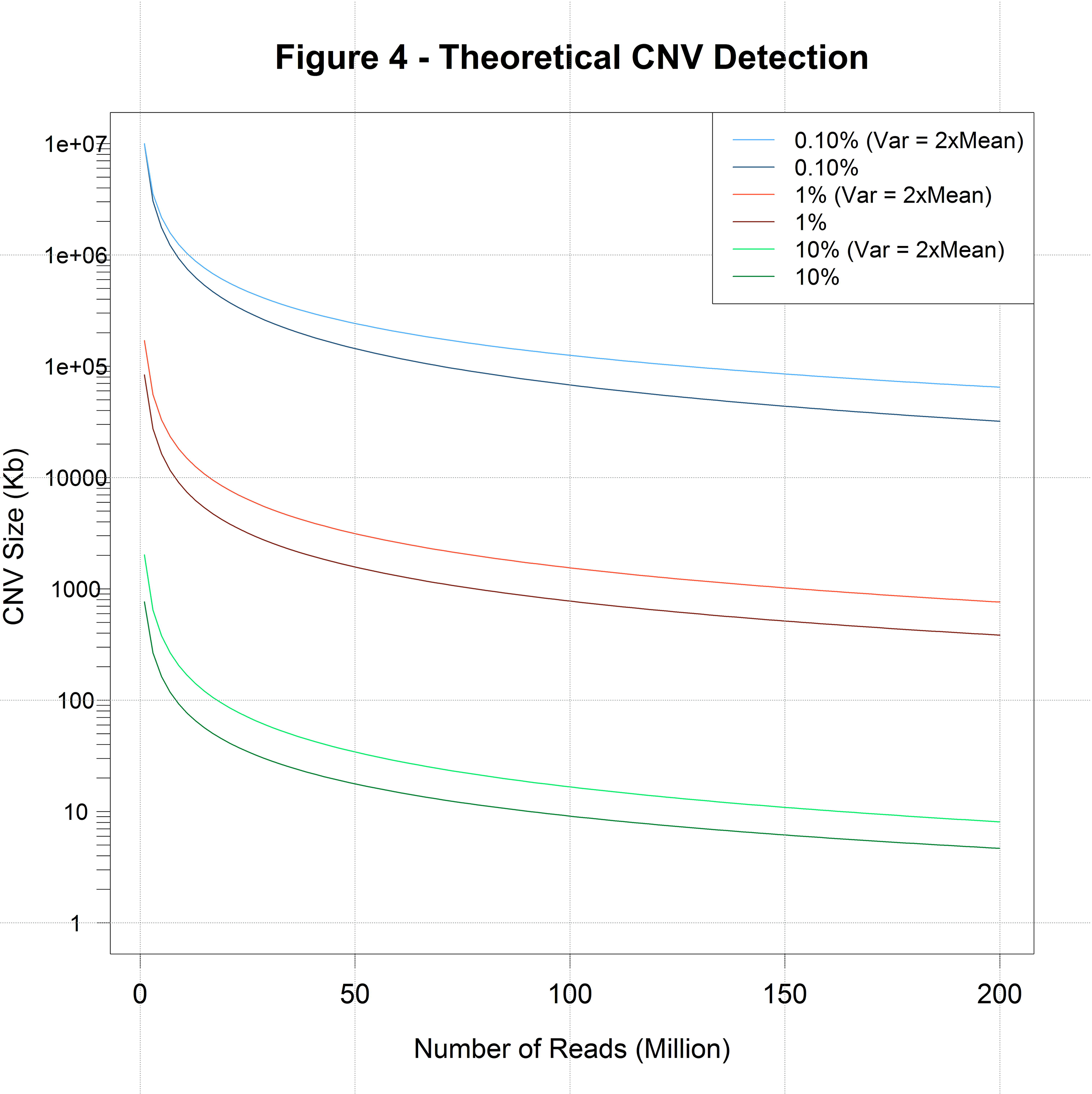
Theoretical power of CNV Detection. CNV size (Kb) detectable with a p-value < 0.01 is plotted against the number of reads required to do so. Each curve represents the amount of ctDNA as a percentage of total circulating DNA. Darker colored curves represent calculations using a Negative Binomial (NB) model and lighter colors represent calculations using a NB model with variance fixed as twice the mean.

## DISCUSSION

Early detection is the key to improving cancer outcomes; however, modern screening methods have false positive rates that expose the healthy population to unacceptable risk. Interest in ctDNA screening methods has grown tremendously given recent reports of cancer detected incidentally during NIPT screening. However, point mutations have been the focus of ctDNA methods developed to date – methods that are extremely useful for monitoring of cancer treatment response (17-19). Yet, for early stage tumors, the fraction of ctDNA amongst the total circulating DNA pool severely limits the practical application of these technologies for cancer screening. At 0.1% ctDNA fraction of total circulating DNA, one can expect only 1 – 5 mutant DNA copies per mL of blood(16) – a mutation fraction that is below the error rate of modern sequencing platforms and detectable only by highly specialized and targeted assays (17). The narrow focus of these assays limits their utility for broad cancer screening, even for a single tumor type, with some exceptions for tumor types like melanoma where a single point mutation (BRAF V600E) is highly recurrent. On the other hand, large tumor-derived copy number aberrations will contribute millions of DNA fragments to the overall circulating DNA pool. The analyses presented herein suggest that it is feasible to detect these large tumor-derived CNVs in circulating DNA with reasonable sequencing depth, and that the detectable CNV profiles can accurately determine the organ of origin of the tumor – a feature that is incredibly important for early stage screening purposes.

While the predictive segments were selected via an unbiased methodology, they demonstrate significant overlap with recurrent CNVs identified in a pan cancer study across 12 common cancer types (24). About 50% of the important segments at the 5 Mb segment size threshold contained a recurrent pan-cancer amplification or deletion and another 30% of the segments overlap with disease specific amplifications or deletions. Moreover, 8 of 10 of the most important segments for cancer type determination were among this list of recurrently amplified and/or deleted pan-cancer CNVs. These most important amplified or deleted segments contain known cancer genes including MYC, EGFR, ERBB2 CDKN2A, RB1 and STK11. For example, 8q24 (25-27) aberrations are suggestive of cancer across a large variety of tumor types.

Interestingly, the tumor types that could be most effectively identified via CNV profile did not necessarily align with the M-class (mutation class) vs. C-class (copy number variant class) tumor types as determined by another pan-cancer study (28). For example, although KIRC is an M-class tumor, it was among the most effectively classified tumor types based on its CNV profile. While KIRC is not broadly copy number aberrant, loss of the short arm of chromosome 3 (3p) (29) containing genes like VHL, PBRM1, BAP1 and SETD2, is highly predictive of KIRC – given it is observed in almost 90% of KIRC cases. Thus, the promise of this approach is not necessarily limited to C-class tumor types.

Some potential challenges for the implementation of ctDNA CNV detection for early cancer screening are not fully addressed by this *in silico* analysis. For example, it will be necessary to understand and override the issue of sample variability in order to achieve the accurate identification of CNVs via ctDNA. While we have made a conservative estimate of this variability, robust sequencing-depth normalization schemes will likely be necessary to achieve this level of variance. Moreover, it is presumed, but not known, whether CNVs predictive of cancer are present in early stage tumors. The battery of tumor profiles used in this analysis, derived from The Cancer Genome Atlas (http://cancergenome.nih.gov/), contain many late stage tumors. These analyses suggest that cancer screening via ctDNA based CNV detection should be attempted in diverse and larger patient cohorts.

## METHODS

### Copy Number Variation Data

We downloaded whole genome copy number variation data, generated by the Tumor Cancer Genome Atlas Research Network (http://cancergenome.nih.gov/), for 25 different cancer types: Adrenocortical carcinoma (ACC), Bladder Urothelial Carcinoma (BLCA), Brain Lower Grade Glioma (LGG), Breast invasive carcinoma (BRCA), Cervical squamous cell carcinoma and endocervical adenocarcinoma (CESC), Colon adenocarcinoma (COAD), Esophageal carcinoma (ESCA), Glioblastoma multiforme (GBM), Head and Neck squamous cell carcinoma (HNSC), Kidney Chromophobe (KICH), Kidney renal clear cell carcinoma (KIRC), Kidney renal papillary cell carcinoma (KIRP), Liver hepatocellular carcinoma (LIHC), Lung adenocarcinoma (LUAD), Lung squamous cell carcinoma (LUSC), Ovarian serous cystadenocarcinoma (OV), Pancreatic adenocarcinoma (PAAD), Pheochromocytoma and Paraganglioma (PCPG), Prostate adenocarcinoma (PRDA), Rectum adenocarcinoma (READ), Skin Cutaneous Melanoma (SKCM), Stomach adenocarcinoma (STAD), Thyroid carcinoma (THCA), Uterine Carcinosarcoma (UCS) and Uterine Corpus Endometrial Carcinoma (UCEC). The data was accessed in December 2014 from *http://gdac.broadinstitute.org/* (30) (doi:10.7908/C19P30S6).

Specifically, we downloaded the segmentation files which contain information about the copy number of segmented genomic data produced by various algorithms like GLAD and CBS(31, 32). Variants in each sample were first run through the SG-ADVISER CNV annotation pipeline (33) and then variants with an allele frequency of >1% in the 1000 Genomes (34) or the Wellderly (35, 36) cohorts were filtered out.

### Data Representation

We adapted the SAX transformation (20) to represent the CNV data in a concise format while not losing any critical information. We first divided chromosomes 1-22 into segments of sizes 5, 6, 7.5, 10, 30, 40, 60, 75, 100 Mega bases. For each of the segments we calculated the average segment duplication value.

This average segment duplication value was then mapped to an appropriate letter representation. Specifically, any value above 0.4 was mapped to ‘**e**’, values between 0.2 and 0.4 were mapped to ‘**d**’, values between −0.2 and 0.2 were mapped to ‘**c**’, values between −0.2 and −0.4 were mapped to ‘**b**’ and any value below −0.4 were mapped to ‘**a**’. We chose a cardinality of 5 to be able to represent normal or 2 copies, 1 copy amplification, more than 2 copy amplification, 1 copy deletion and 2 copy deletion of the genomic segment.

### Prediction Methods

The SAX transformed data was used to train machine learning models which could distinguish a normal sample from a cancer sample and also predict the type of the cancer as described by TCGA. We used random forests which is an ensemble learning method and the KNN (k-nearest neighbors) method which is a simpler pattern recognition algorithm.

The standard k-Nearest Neighbor algorithm was implemented using custom code in R. Briefly, we calculated distances between each of the SAX transformed samples using a modified hamming distance metric where the distance between adjacent letters was fixed as 0.5 and any other changes were fixed as 1. The data was then randomly split into a training set (75%) and a test set (25%). For classification the custom distance metric was used to find k training samples closest to the test sample and the majority class in the k samples was assigned to the test sample. k was set as the square root of the total number of samples.

A separate random forest model was trained to optimize overall accuracy for each segment size threshold using a 10 fold cross validation training scheme, each model contained 100 trees and the optimal number of variables randomly sampled as candidates at each split in the trees was determined heuristically.

For tumor clustering we used the segments marked as important by the random forest model. Then we used the modified hamming distance described earlier to calculate the distances for the clustering. Each column in the heat map corresponds to a sample, and the rows represent the important segments identified by the random forest. Each cell has been colored red for a gain and blue for a loss.

For the tumor misclassification heat maps the color scale was used to represent the percentage of misclassification as another tumor type or normal with white being 0% and dark blue being 100% of the misclassifications. The samples were clustered according to a custom similarity metric S. Let A and B be two tumor samples with A_n_ and B_n_ being the number of samples in A and B respectively, then if α and β are the fraction of samples of A classified as B and the fraction of samples of B classified as A, and N is the total number of samples, the similarity S(A,B) between A and B can be defined as -

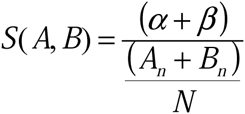

### Theoretical Power of CNV Detection Calculations

To calculate the theoretical limit of detecting CNVs we used a negative binomial distribution to model the sequencing of circulating DNA and subsequent read mapping. Generally, sequencing data is affected by biases in genome composition, sequencing and mapping and thus a negative binomial distribution does a better job at modelling the sample variance as compared to a Poisson distribution (37). We used two separate negative binomial models, for the first we used the following definition of the probability mass function (pmf)

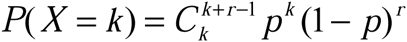

Where X is the random variable denoting the number of successes before ‘r failures’ or in this case number of reads mapped to the genomic segment of interest; p is the probability of one success i.e. the probability of mapping a read to that genomic segment calculated as 1/(Number of Segments); r, the number of failures, is calculated as (1-p)(Total number of reads) denoting the number of reads mapped to any other genomic region. For the second model we used an alternative formulation of the negative binomial distribution represented as the following pmf

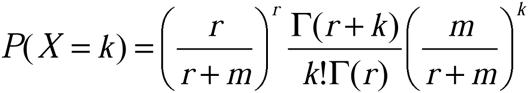

In this case r is referred to as the “dispersion parameter” or the “shape parameter” and m is the mean of the distribution calculated as (Total number of reads)/(Number of segments). The variance for this model is given by 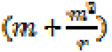. We fixed the variance as twice the mean value, thus getting an estimate for ‘r’ and using that for the model. This was done in order to simulate an arbitrarily large variance. For both these models we then calculated the right tail of the pmf for X being equal to the expected number of counts at a segment in case of a 1 copy duplication event given the mean being equal to the expected counts at the segment in case of a normal sample, thus getting a p-value. We performed these calculations at various read depths and circulating tumor DNA representing 0.1%, 1% and 10% fraction of the total circulating DNA in the sample.

### Software

Data filtration and SAX transformations were performed using custom scripts in python. All models were built using R v3.1.1. The ‘caret’ library in R was used to train the random forest models. ROC curves denoting the performance of the models were plotted using the library ‘pROC. Heat maps were plotted using the ‘gplots’ library. All calculations for the theoretical limit of CNV detection were performed using R version 3.1.1 as well.

## ACKNOWLEDGMENTS

This work is supported by a Scripps Translational Science Institute pilot award (5 UL1 TR001114) from Scripps Genomic Medicine, a NIH-NCATS Clinical and Translational Science Award (CTSA; 5 UL1 RR025774) to STSI. Further support is provided by NHGRI Genome Sequencing Informatics Tools (GS-IT) Program via grant NIH U01 HG006476.

## DISCLOSURE DECLARATION

Ali Torkamani declares he is a co-founder of Cypher Genomics, Inc. Bhuvan Molparia declares no conflict of interest. Eshaan Nichani declares no conflict of interest.

## REFERENCES

1. Suh KS, Park SW, Castro A, Patel H, Blake P, Liang M, et al. Ovarian cancer biomarkers for molecular biosensors and translational medicine. Expert review of molecular diagnostics. 2010;10(8):1069–83.

2. Menon U, Gentry-Maharaj A, Hallett R, Ryan A, Burnell M, Sharma A, et al. Sensitivity and specificity of multimodal and ultrasound screening for ovarian cancer, and stage distribution of detected cancers: results of the prevalence screen of the UK Collaborative Trial of Ovarian Cancer Screening (UKCTOCS). The Lancet Oncology. 2009;10(4):327–40.

3. Hubbard RA, Kerlikowske K, Flowers CI, Yankaskas BC, Zhu W, Miglioretti DL. Cumulative probability of false-positive recall or biopsy recommendation after 10 years of screening mammography: a cohort study. Annals of internal medicine. 2011;155(8):481–92.

4. Imperiale TF, Ransohoff DF, Itzkowitz SH, Levin TR, Lavin P, Lidgard GP, et al. Multitarget stool DNA testing for colorectal-cancer screening. The New England journal of medicine. 2014;370(14):1287–97.

5. Punglia RS, D’Amico AV, Catalona WJ, Roehl KA, Kuntz KM. Effect of verification bias on screening for prostate cancer by measurement of prostate-specific antigen. The New England journal of medicine. 2003;349(4):335–42.

6. Buys SS, Partridge E, Black A, Johnson CC, Lamerato L, Isaacs C, et al. Effect of screening on ovarian cancer mortality: the Prostate, Lung, Colorectal and Ovarian (PLCO) Cancer Screening Randomized Controlled Trial. Jama. 2011;305(22):2295–303.

7. Shlien A, Malkin D. Copy number variations and cancer. Genome medicine. 2009;1(6):62.

8. Kandoth C, McLellan MD, Vandin F, Ye K, Niu B, Lu C, et al. Mutational landscape and significance across 12 major cancer types. Nature. 2013;502(7471):333–9.

9. Carrasco DR, Tonon G, Huang Y, Zhang Y, Sinha R, Feng B, et al. High-resolution genomic profiles define distinct clinico-pathogenetic subgroups of multiple myeloma patients. Cancer Cell. 2006;9(4):313–25.

10. Hoadley KA, Yau C, Wolf DM, Cherniack AD, Tamborero D, Ng S, et al. Multiplatform analysis of 12 cancer types reveals molecular classification within and across tissues of origin. Cell. 2014;158(4):929–44.

11. Hofree M, Shen JP, Carter H, Gross A, Ideker T. Network-based stratification of tumor mutations. Nature methods. 2013;10(11):1108–15.

12. Heitzer E, Auer M, Hoffmann EM, Pichler M, Gasch C, Ulz P, et al. Establishment of tumor-specific copy number alterations from plasma DNA of patients with cancer. International journal of cancer Journal international du cancer. 2013;133(2):346–56.

13. Hughes V. Pregnant Women Are Finding Out They Have Cancer From A Genetic Test Of Their Babies: BuzzFeed News; 2015. Available from: http://www.buzzfeed.com/virginiahughes/pregnant-women-are-finding-out-they-have-cancer-from-a-genet#.ncjvA5VNz8.

14. Nygren AOH, Dean J, Jensen TJ, Kruse S, Kwong W, van den Boom D, et al. Quantification of Fetal DNA by Use of Methylation-Based DNA Discrimination. Clinical Chemistry. 2010;56(10):1627–35.

15. Fan HC, Gu W, Wang J, Blumenfeld YJ, El-Sayed YY, Quake SR. Non-invasive prenatal measurement of the fetal genome. Nature. 2012;487(7407):320–4.

16. Dawson SJ, Tsui DW, Murtaza M, Biggs H, Rueda OM, Chin SF, et al. Analysis of circulating tumor DNA to monitor metastatic breast cancer. The New England journal of medicine. 2013;368(13):1199–209.

17. Newman AM, Bratman SV, To J, Wynne JF, Eclov NC, Modlin LA, et al. An ultrasensitive method for quantitating circulating tumor DNA with broad patient coverage. Nature medicine. 2014;20(5):548–54.

18. Zill OA, Greene C, Sebisanovic D, Siew LM, Leng J, Vu M, et al. Cell-Free DNA Next-Generation Sequencing in Pancreatobiliary Carcinomas. Cancer Discov. 2015.

19. Crowley E, Di Nicolantonio F, Loupakis F, Bardelli A. Liquid biopsy: monitoring cancer-genetics in the blood. Nat Rev Clin Oncol. 2013;10(8):472–84.

20. Lin J, Keogh E, Lonardi S, Chiu B. A Symbolic Representation of Time Series, with Implications for Streaming Algorithms. In proceedings of the 8th ACM SIGMOD Workshop on Research Issues in Data Mining and Knowledge Discovery. 2003.

21. Chan KC, Jiang P, Zheng YW, Liao GJ, Sun H, Wong J, et al. Cancer genome scanning in plasma: detection of tumor-associated copy number aberrations, single-nucleotide variants, and tumoral heterogeneity by massively parallel sequencing. Clin Chem. 2013;59(1):211–24.

22. Leary RJ, Sausen M, Kinde I, Papadopoulos N, Carpten JD, Craig D, et al. Detection of chromosomal alterations in the circulation of cancer patients with whole-genome sequencing. Science translational medicine. 2012;4(162):162ra54.

23. Fernández-Delgado M, Cernadas E, Barro S, Amorim D. Do we Need Hundreds of Classifiers to Solve Real World Classification Problems?. Journal of Machine Learning Research. 2014;15:3133–81.

24. Zack TI, Schumacher SE, Carter SL, Cherniack AD, Saksena G, Tabak B, et al. Pan-cancer patterns of somatic copy number alteration. Nature genetics. 2013;45(10):1134–40.

25. Witte JS. Multiple prostate cancer risk variants on 8q24. Nature genetics. 2007;39(5):579–80.

26. Salinas CA, Kwon E, Carlson CS, Koopmeiners JS, Feng Z, Karyadi DM, et al. Multiple independent genetic variants in the 8q24 region are associated with prostate cancer risk. Cancer epidemiology, biomarkers & prevention: a publication of the American Association for Cancer Research, cosponsored by the American Society of Preventive Oncology. 2008;17(5):1203–13.

27. Jia L, Landan G, Pomerantz M, Jaschek R, Herman P, Reich D, et al. Functional enhancers at the gene-poor 8q24 cancer-linked locus. PLoS genetics. 2009;5(8):e1000597.

28. Ciriello G, Miller ML, Aksoy BA, Senbabaoglu Y, Schultz N, Sander C. Emerging landscape of oncogenic signatures across human cancers. Nature genetics. 2013;45(10):1127–33.

29. Zbar B, Brauch H, Talmadge C, Linehan M. Loss of alleles of loci on the short arm of chromosome 3 in renal cell carcinoma. Nature. 1987;327(6124):721–4.

30. Analysis-ready standardized TCGA data from Broad GDAC Firehose stddata__2015_02_04 run. In: Institute B, editor. 2015.

31. Hupe P, Stransky N, Thiery JP, Radvanyi F, Barillot E. Analysis of array CGH data: from signal ratio to gain and loss of DNA regions. Bioinformatics. 2004;20(18):3413–22.

32. Olshen AB, Venkatraman ES, Lucito R, Wigler M. Circular binary segmentation for the analysis of array-based DNA copy number data. Biostatistics. 2004;5(4):557–72.

33. Erikson GA, Deshpande N, Kesavan BG, Torkamani A. SG-ADVISER CNV: copy-number variant annotation and interpretation. Genetics in medicine: official journal of the American College of Medical Genetics. 2014.

34. Genomes Project C, Abecasis GR, Auton A, Brooks LD, DePristo MA, Durbin RM, et al. An integrated map of genetic variation from 1,092 human genomes. Nature. 2012;491(7422):56–65.

35. Borrell B. Sequencing projects bring age-old wisdom to genomics. Nature medicine. 2011;17(11):1329.

36. Singleton MV, Guthery SL, Voelkerding KV, Chen K, Kennedy B, Margraf RL, et al. Phevor combines multiple biomedical ontologies for accurate identification of disease-causing alleles in single individuals and small nuclear families. American journal of human genetics. 2014;94(4):599–610.

37. Sampson J, Jacobs K, Yeager M, Chanock S, Chatterjee N. Efficient study design for next generation sequencing. Genetic epidemiology. 2011;35(4):269–77.

